# A new subspecies of *Ficus* L. (Moraceae) from North Andaman Island, India

**DOI:** 10.1101/2023.10.16.562493

**Authors:** Raja Kullayiswamy Kusom, SarojiniDevi Naidu

## Abstract

*Ficus pellucidopunctata* subsp. *obpyriformis* is described and illustrated from the North Andaman Island of India. The new taxon resembles *Ficus pellucidopunctata* in punctale leaf, leaf shape and venation, but differs with the typical species by its longer and obpyriform fig, (verses ellipsoid); scaly minute (ca. 2 mm) basal bracts that do not cover the base of the fig (verses larger (ca. 6 mm) basal bracts that cover the base of the fig); fig orange to dark brown (verses green to yellow); branches terete (verses. angular). *Ficus pellucidopunctata* subsp. *obpyriformis* seeds germinated and saplings were planted and are growing in the Dharmavana Nature Ark (DNA), Bhongir, Telangana as part of the DNA’s *exsitu* conservation programme.

## Introduction

The genus *Ficus* L. Moraceae is commonly known as ‘Fig’ due to the fruits which are a special type of inflorescence called ‘hypanthodium’. This cup-shaped receptacle is formed by the condensation of the rachis into densely packed cymes, wherein male, female and gall flowers are arranged in groups. *Ficus* also is known for its well diversified traits such as deciduous and evergreen trees, shrubs, climbers and creepers, and life forms such as epiphytes, semi-epiphytes growing in crevices, rheophytes, and lithophytes (Sudhakar et al, 2017).

*Ficus* species are largely distributed in tropical and subtropical forests where many serve as keystone species within the ecosystem. Their figs attract insects, birds and other animals by giving food throughout the year (Chaudhary et al, 2012; Corner, 1965, Berg, 1989; Berg and Corner, 2005). *Ficus* is one of the largest genera of angiosperms with around 750 species. Sixty-six percent of the world’s fig species are concentrated in the Asian-Australian region of about 500 species (Chaudhary et al, 2012).

King (1887-1888) worked on British India *Ficus* and at that time he recorded 113 species of which 75 are located within the present political boundaries of India. Following King’s effort, much exploration and research has been conducted throughout the world (Corner, 1958, 1965, 1981; Berg, 1989, 2003, 2004 a, b; Dixon, 2003; Burrows & Burrows, 2003; Berg and Corner, 2005; Chaudhary *et al*., 2012). These efforts have brought considerable changes in identification, nomenclature and distributional patterns. No dedicated account for species distributed throughout modern India existed till Chaudary et al. (2012) made a list of 115 taxa of which 10 are endemic. A schematic representation of the Chaudary *et al*. (2012) article has been provided in this article for a better understanding of genus *Ficus* classification.

While on travel to visit a friend in Diglipur, North Andaman, Mr. Dürr noticed a *Ficus* tree on the roadside with which he was unacquainted. He photographed the tree and subsequently collected figs and foliage which he presented to the authors for identification. After critical examination of images and samples, the authors prepared herbarium specimens. The collected material was examined carefully by the authors, comparing with existing species in regional and local floras, and other herbarium specimens along with online databases. The authors concluded that the collected specimens came closest to *Ficus pellucidopunctata* Griff. They then compared both for differentiating morphological characters. Tabulated phenological characters in Table 1 show variations mainly in ‘synconia’, structure, shape and size that are adequate and considerable to establish a new subspecies under the type species. Given those variations the authors raised a new subspecies under *F. pellucidopunctata* as classified under the genus *Ficus*, subspecies *Urostigma*, and section *Urostigma*, subsection *Conosycea. Ficus pellucidopunctata* is distributed throughout Indochina into Peninsular Malaysia, Borneo and the Philippines but nowhere common.

**Table 1:**
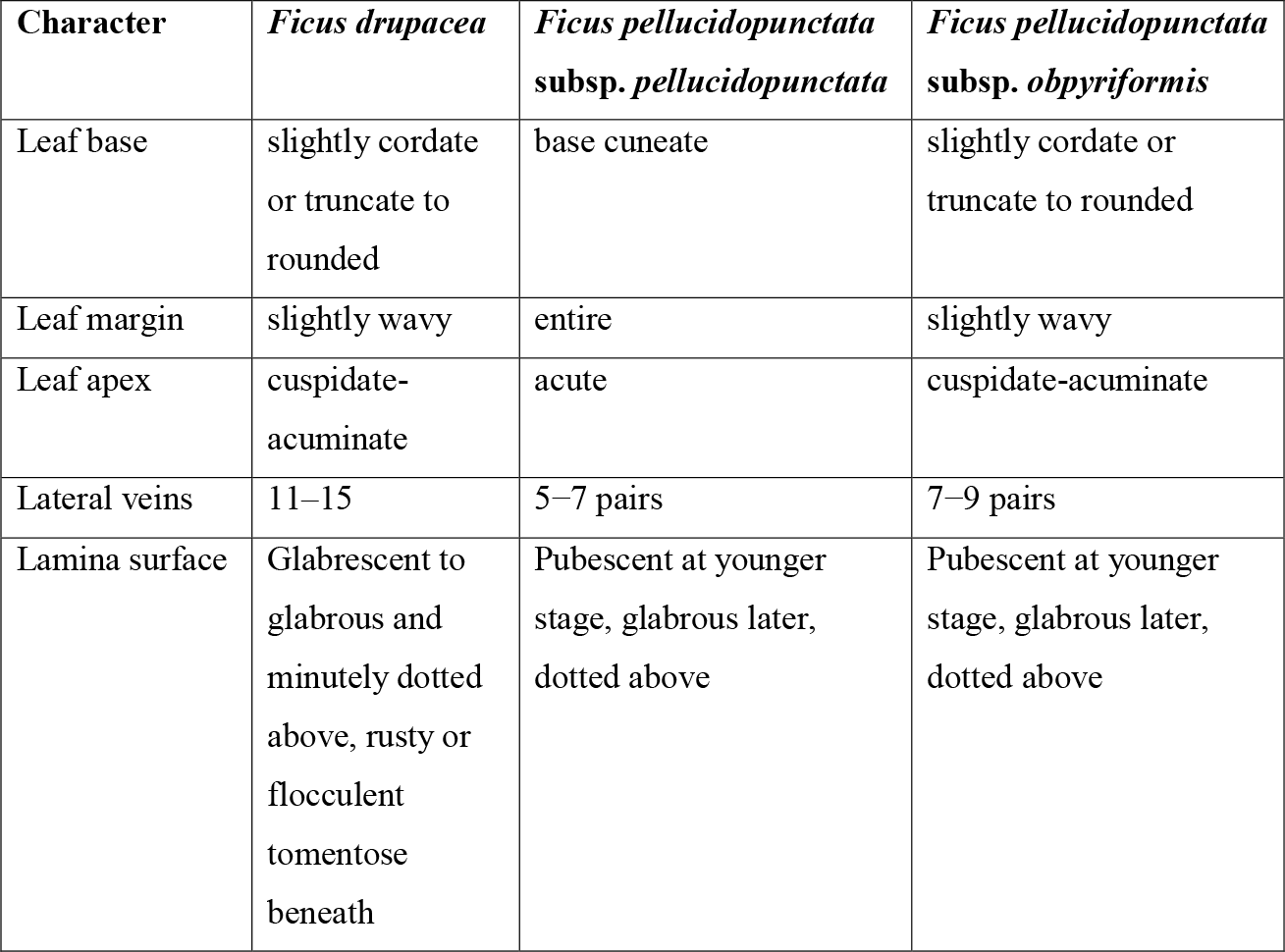
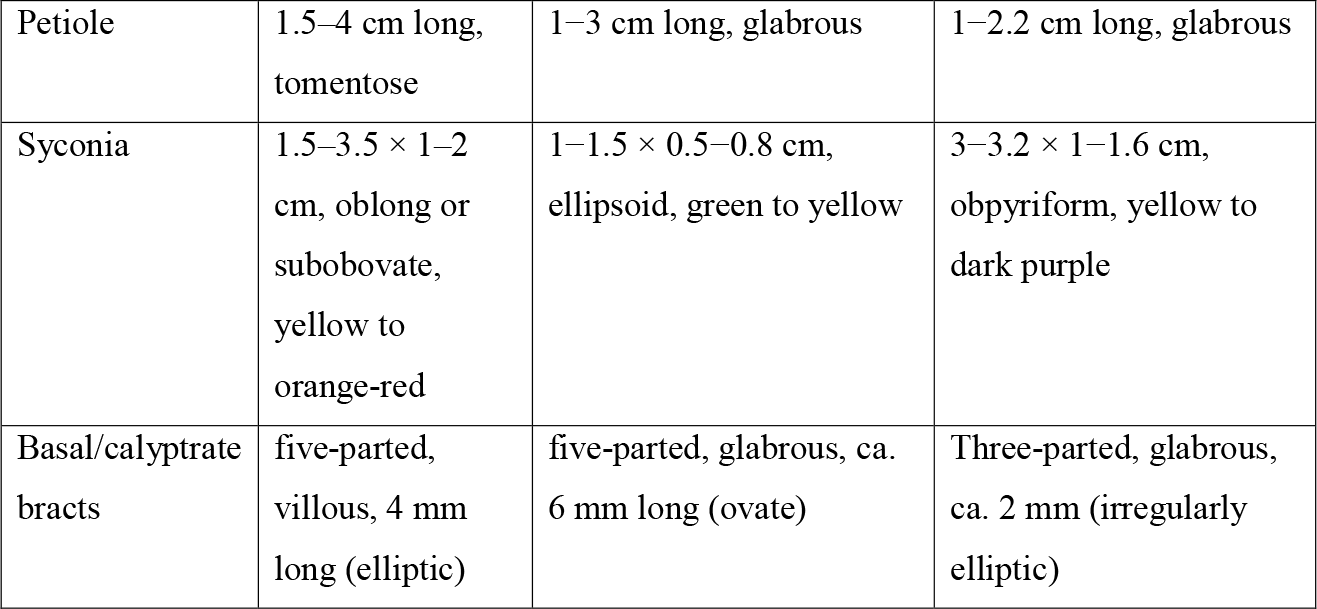
Differences between *Ficus drupacea*, typical species *Ficus pellucidopunctata*, and the new subsp. *obpyriformis*.

## Materials & Methods

Collected specimens were treated with ethanol and herbarium specimens were prepared following the standard methods (Santapau, 1958, Jain & Rao, 1977). Figs were used for microscopic observations and imaging, and two figs were preserved in 70% ethanol for future reference. Microscopic images were taken with an Olympus stereo microscope SZ61, Magcam DC5 camera attached. Seeds of the same were germinated for *ex-situ* conservation in the Dharmavana Nature Ark (DNA), Yadadri-Bhuvanagiri, Telangana.

## Taxonomic treatment

***Ficus pellucidopunctata* subsp. *obpyriformis* subspecies nova** (**Fig. 1 & 2**)

**Fig 1:**
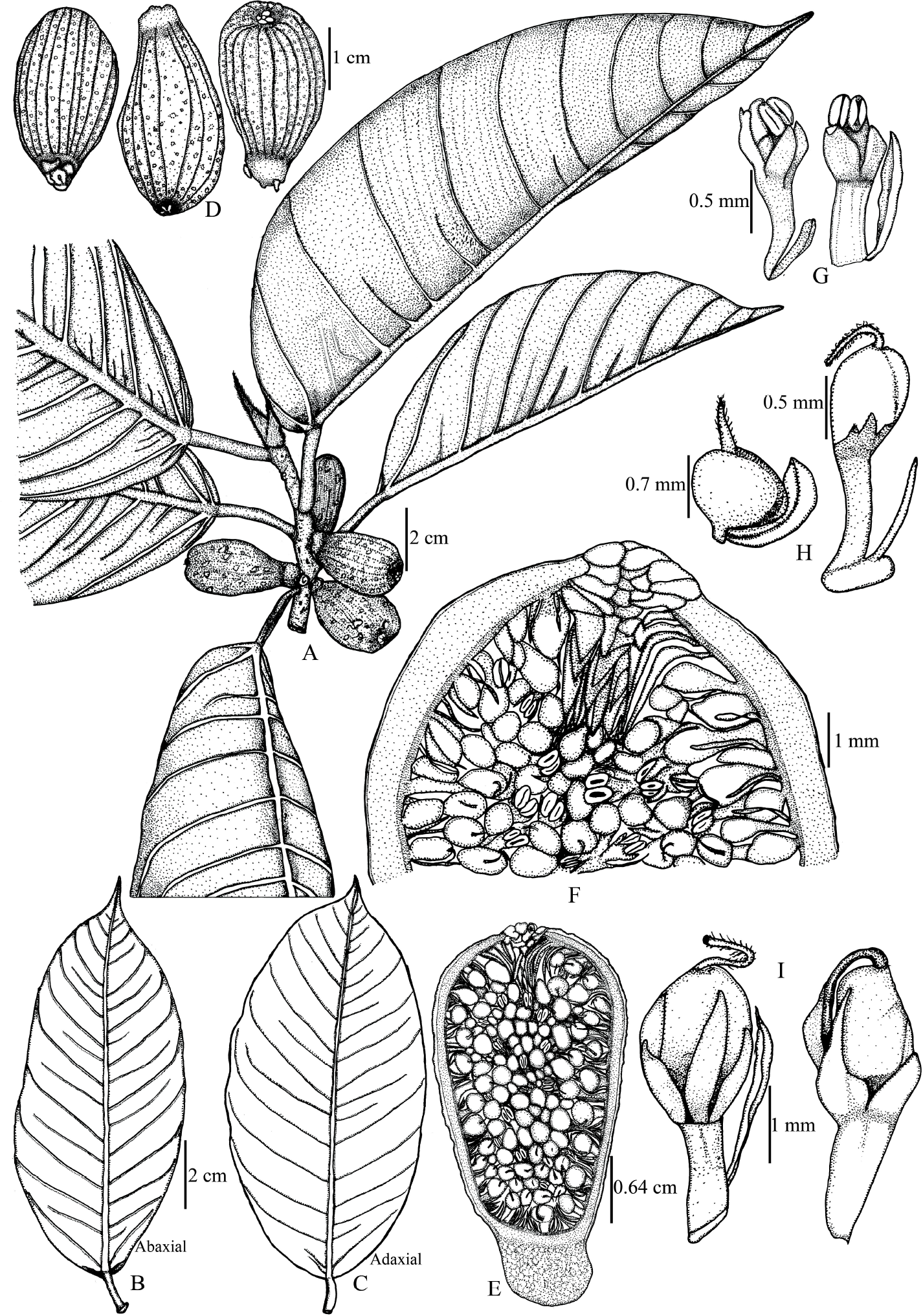
*Ficus pellucidopunctata* subsp. *obpyriformis* subspecies nova: **A**. Twig, **B & C**. Leaves, **D**. Figs, **E**. L.S of Fig, **F**. Closer view of ostiole and flowers in fig, **G**. Male flowers, **H**. Gall flowers, **I**. Female flowers.

**Fig 2:**
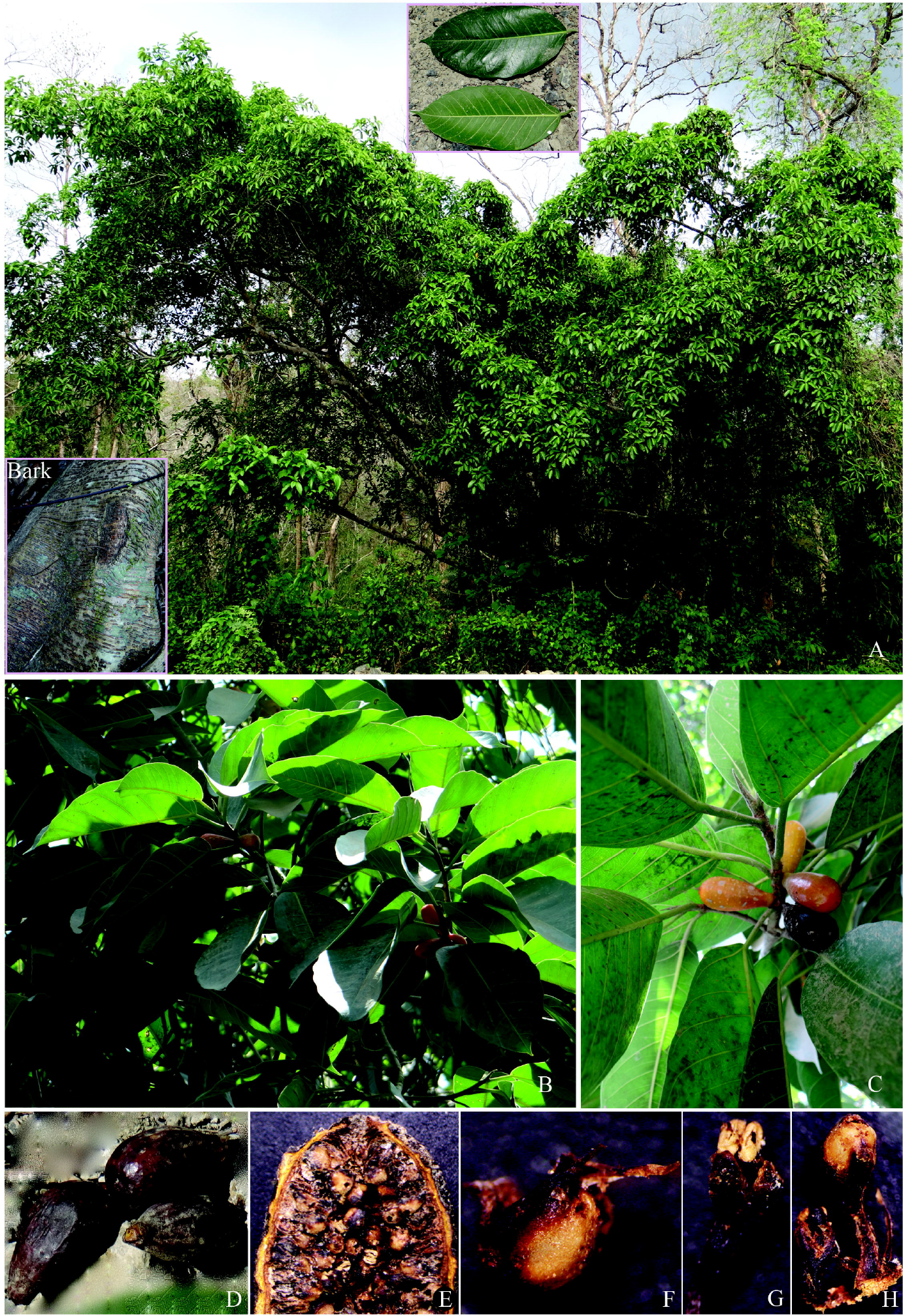
**A**. Habit, **B**. Figs at younger stage, **C**. Fruiting branch, **D**. Figs at mature stage, **E**. Fig L.S, **F**. Gall flower, **G**. Male flower, **H**. Female flower.

Type: India, Andaman and Nicobar Islands, Andamans district, Diglipur Tehsil, near Nischintapur (RV) village, Great Andaman Trunk Road, 13°11’22.0”N 92°52’50.2”E, 71 m, *W. F. Dürr* DNA196 (Holotype BSID!; Isotypes MH!, HDNA!).

This new taxon differs with the typical *Ficus pellucidopunctata* by its longer and obpyriform fig, (vs. ellipsoid); scaly minute (ca. 2 mm) basal bracts that not cover the base of the fig (vs. larger (ca. 6 mm) basal bracts that cover base of the fig); fig orange to dark brown (vs. green to yellow); branches terete (vs. angular).

***Note*:** Only one individual of this new taxon was found in the type location. The DNA is growing many individuals from seed and conserving this taxon at the DNA site. Nearly all other *Ficus* species of India are conserved in the Dharmavana Nature Ark. Photographs of seedlings are included in this article. At first, the seedlings show a remarkable resemblance to those of *Ficus drupacea* (Fig. 3 A&B).

**Fig 3:**
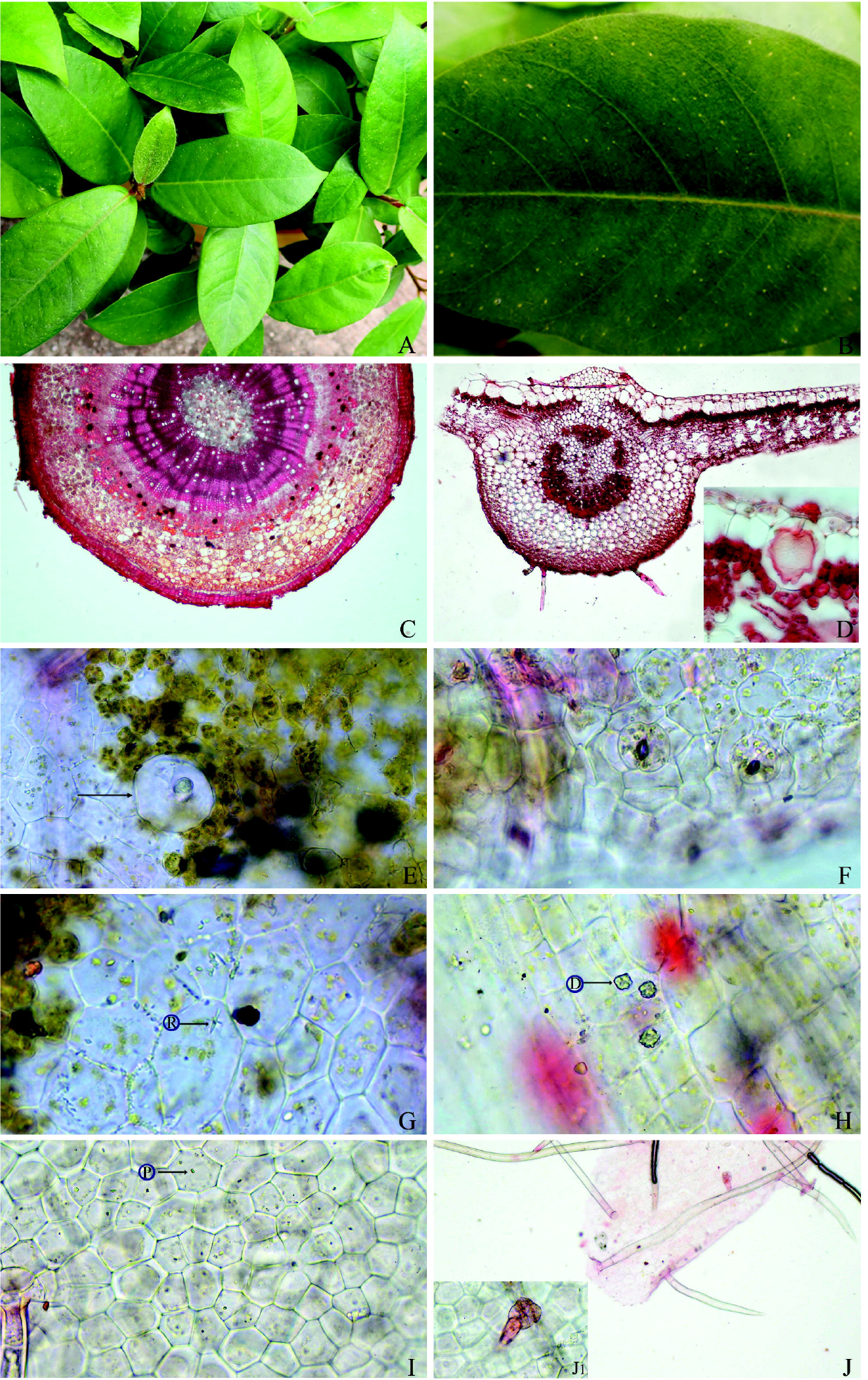
**A**. Seedling, **B**. Leaf showing punctate glands, **C**. T.S of stem, **D**. T.S of leaf along with cystolith, E-G Leaf adaxial: **E**. Punctate gland on upper-surface along with crystals, **F**. Stomata (Anomocytic), **G**. Star-shaped raphid, H-J Leaf abaxial: **H**. Stellate drusa, **I**. Lower epidermis with prism-shaped crystals, **J**. Glandular and non-glandular trichomes, **J1**. Glandular trichome enlarged.

### Description

Tree of about 15 m height with few aerial roots, bark grey and green with horizontal tuberculate, branches spreading, terete, fibres, glabrous when young, rough when mature. Leaf alternate, rigid, flat, early pubescent later glabrous, dark brown when dry; petiole canaliculated, glabrous, 1−2.2 cm long, 2−3.5 mm thick; lamina ovate, elliptic, elliptic-oblong, 7.5−17 × 3−8.2 cm, base slightly cordate or truncate or rounded to tapering, margin entire, slightly wavy, apex cuspidate-acuminate (up to 1.5 cm long), mid-nerve very prominent below, slight above, lateral 7−9 pairs of primary veins looped, and 15−19 pairs of secondary veins conspicuous, in between primary veins are arcuate; bud scale lanceolate, 1− 2 cm long, 1 cm wide, villous outside, glabrous inside, brown, sharply acuminate. Basal bracts of fig 3, irregularly elliptic, 0.5 × 0.7 cm, boat-shaped, light brown. Figs axillary, sessile, glabrous, smooth, shiny, slightly furrowed, in pairs, obpyriform, 3−3.2 × 1− 1.6 cm; base 0.5 × 0.7 cm, cylindrical; apex rounded to truncate, 0.8 cm wide (with a rosette) composed of small ostiolar bracts; fig-wall slightly thick, 1.5−2 mm; ostiolar bracts 3, triangular, 1−2 × 2 mm; internal ostiolar bracts rather longer than the flower up to 4 mm long, dark purple, glabrous. Male flowers 1.5 mm long, disperse all along, frequent, sub-sessile, bracts 3, ovate, 1 × 0.5 mm, stamens 2, white, 0.52 mm long, 0.72 mm wide two together, dehiscence longitudinally; sessile gall flower 1.2 mm long including style part, bract as long as gall flower, boat-shaped, brown, 3 nerved; pedicellate gall flower 1.5 mm long, perianth as sessile gall flower; female flowers 3 mm long, pedicel 0.8−1.5 mm, bracts brown 1.5 mm long, glabrous, thine; perianth lobes 1–2, triangular to ovate, brown with papery transparent margin, stigma simple, slightly bristle. Seed slightly keeled, light brown (whitish), subglobose, 1.5 mm dia. Cystoliths subglobose, potruberent, stomata anomocytic (190 µm across), present on adaxial only, guard cell 50 × 70 µm; calcium oxalate crystals present on both adaxial and abaxial, they are raphids (R), druses (D), prisms (P).

### Flowering & Fruiting

- January – April.

### Habit and Habitat

The new taxon grows up to 15 metres high in black soil of semi-deciduous forest. The species associated with the *Ficus pellucidopunctata* subsp. *obpyriformis* are the canopy species *Ficus virens, F. nervosa, F. hispida, Holoptelia integrefolia, Butea monosperma, Terminalia bialata, Lagerstroemia sp*., *Dalbergia sp*., *Albizia sp*., *Mytragyna parvifolia* and under canopy species *Calamus nagbettai, Musa* sp., *Tylophora* sp. *Passiflora* sp., *Alpinia* sp. *Pergularia daemia*. The area is being converted for cultivation and forest trees are being logged.

### Etymology

The subspecies epithet *‘obpyriformis’* describes the shape of the syconia.

## Acknowledgements

The authors are grateful to the Dharmavana Nature Ark (DNA) for providing the facilities and support necessary for the completion of this article and to Mr. Dürr for bringing this taxon to authors’ attention.

## Classification diagram

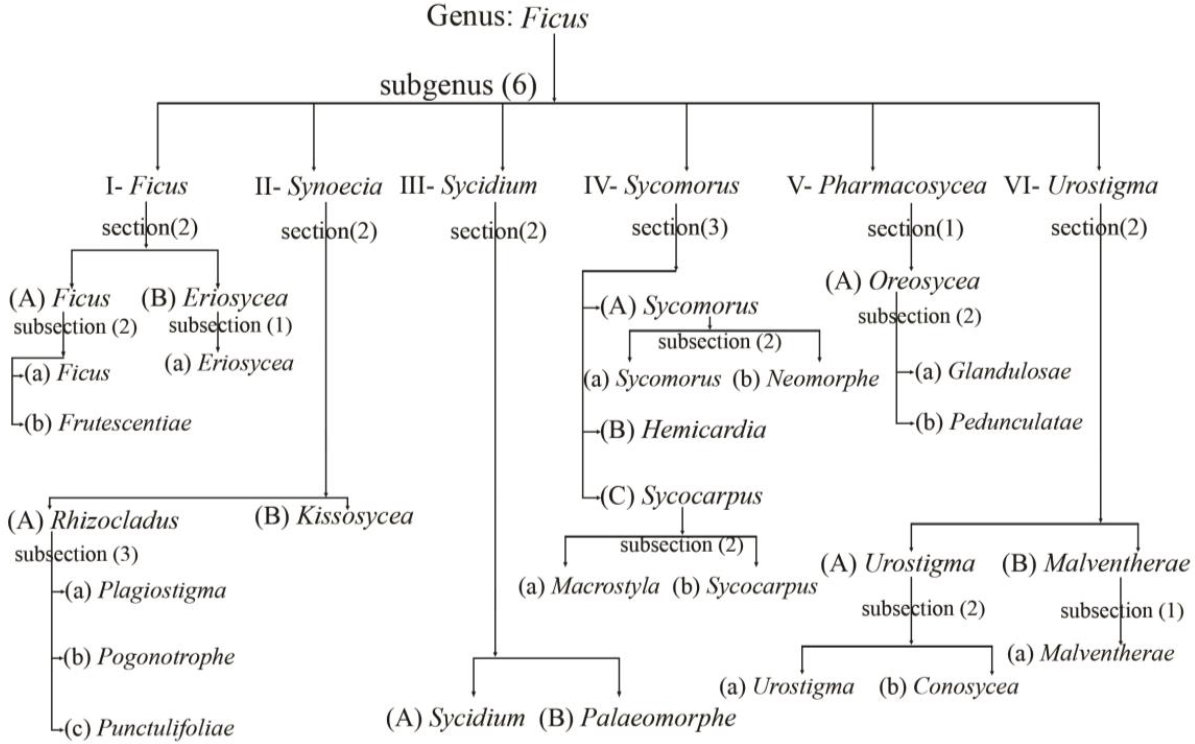

